# Protecting seabed sediment carbon for climate mitigation: a UK case study

**DOI:** 10.1101/2022.02.10.479679

**Authors:** Graham Epstein, Callum M. Roberts

## Abstract

Protecting areas of seabed sediment would appear to be an important climate mitigation strategy, however carbon has not been considered in national management plans. Using spatial analyses we show, in the UK Exclusive Economic Zone (EEZ), high potential organic carbon loss due to mobile bottom fishing is likely to be geographically restricted. Currently designated seabed marine protected areas (MPAs) cover 32.9% of the EEZ, 30.0% of carbon stocks, but only 12.9% of fishing disturbance. If all MPAs were closed to mobile bottom fishing, we estimate a prevention of only 6.9% of potential organic carbon losses due to fishing disturbance across the EEZ. In contrast, areas identified as having largest climate change mitigation potential cover just 16.5% of the EEZ, contain 24.7% of OC stocks and closing these areas to mobile fishing could save 29.6% of potential carbon losses. We must re-evaluate current seabed management plans and incorporate evidence-based carbon considerations.

**Teaser:** Restrictions on bottom trawling and dredging outside of MPA networks may be necessary to derive a climate mitigation benefit.

## 1. Introduction

The ocean has absorbed ∼40% of anthropogenic CO_2_ emissions since the industrial revolution acting as a major brake on climate change (*1, 2*). However, under all future predicted climate scenarios the oceanic carbon sink is projected to become less effective at absorbing CO_2_ from the atmosphere (*3*). One way to improve the ocean’s ability to absorb excess atmospheric CO_2_ is to conserve marine organic carbon (OC) stores and promote natural carbon sequestration (*4*). This process, generally referred to as blue carbon, is well established for marine vegetated habitats (*5*), but it is the appropriate functioning of the entire marine ecosystem which maintains the ocean as a sink for anthropogenic CO_2_ (*6*).

Subtidal marine sediments contain the ocean’s biggest organic carbon store, estimated to hold ∼2.3 Tt in the top 1 m (*7*) and accumulate an additional 126 - 350 Mt of organic carbon per year (*8-10*). However, it is increasingly apparent this store is vulnerable to remobilisation and mineralisation by human disturbance (*8, 11-13*). By far the most widespread human disturbance to the seabed is the use of mobile bottom trawls and dredges to catch fish and shellfish (*14-16*). A global first-order estimate suggested that annually, mobile bottom fishing activities may cause ∼590-1470 Mt of CO_2_ to be released into the water column due to disturbance and remineralisation of organic carbon from seabed sediments (*11*). This is equivalent to 15-20% of the atmospheric CO_2_ absorbed by the ocean each year (*11*). By significantly increasing inorganic carbon concentrations in the ocean, it would slow the rate of CO_2_ uptake from the atmosphere, and possibly release more oceanic CO_2_ back to the atmosphere (*11, 17-19*). Protecting seabed sediment carbon from human disturbance would appear to be an important step in global efforts to mitigate climate change.

Seabed protection is predominantly undertaken through the establishment of marine protected area (MPA) networks (*20*). Such networks have, to date, mainly been designed for biodiversity conservation, emphasising inclusion of declining, depleted, rare or fragile species (*21, 22*). More recently, the potential benefits of MPAs to both mitigate against and adapt to climate change are starting to be recognised, and the protection of carbon stores is being added to the list of conservation objectives (*23, 24*). Although MPAs are now widespread, with ∼8% of the global ocean under a protected area designation (*20*), their conservation objectives and legislation are often limited in scope and therefore many damaging activities still persist (*25*). In total, only around 3% of the global ocean is fully or highly protected from fishing impacts (*20*).

One management option would be to increase the level of protection given to current MPAs to derive a climate mitigation benefit from additional protection of seabed organic carbon stocks. However, as the location of existing MPAs was determined principally by biodiversity and not organic carbon, it is unclear how much benefit this would produce. Additionally, due to historic damage by marine industries, the locations rich in remaining biodiversity, are often distinct from areas of high human activity (*22, 26, 27*). MPAs are therefore frequently located in areas with low human disturbance, offering limited additionality of benefits from protection (*28*). Added benefits of protection of seabed organic carbon for climate mitigation will be greater in place where carbon stores are large and subject to damaging activities.

A common critique of area-based management is that it displaces rather than reduces damaging human activities (*29-31*). Although not ubiquitous (*30*), displacement must be considered when determining climate mitigation potential (*28*). If additional protection of the seabed leads to equivalent losses of organic carbon elsewhere due to displaced human activities then no climate mitigation is achieved (*28*). It is therefore vital to consider and quantify how fisheries displacement may affect organic carbon stocks when designing a protection strategy.

Here, we use the UK exclusive economic zone (EEZ) as a case study to identify areas where protection from bottom fishing activities could have significant climate change mitigation potential. The UK has established one of the most extensive networks of MPAs in the world to protect marine life, consisting of 372 MPAs covering 38% of its EEZ (*32*). Using data on seabed sediment carbon stocks and fishing disturbance, combined with published modelled estimates on the impact of mobile bottom fishing on sediment organic carbon concentrations, we ask is the distribution of potential carbon losses from mobile bottom fishing spatially restricted, how much carbon could be protected by existing MPAs and are additional protected areas, or legislative instruments needed to increase climate change mitigation potential?

## 2. Results

### 2.1 Organic carbon stock and spatial distribution

The study region covered ∼722,697 km^2^ of the UK EEZ, with the northern-most section of the EEZ excluded due to lack of carbon stock data and the most south-westerly section excluded due to lack of fishing activity data (Fig. 1a). By unifying four previously published spatial organic carbon datasets (*33-36*), the study region was estimated to hold 307.2 Mt of organic carbon in the top 10 cm of the seabed (Fig. 1a).

**Fig. 1.**
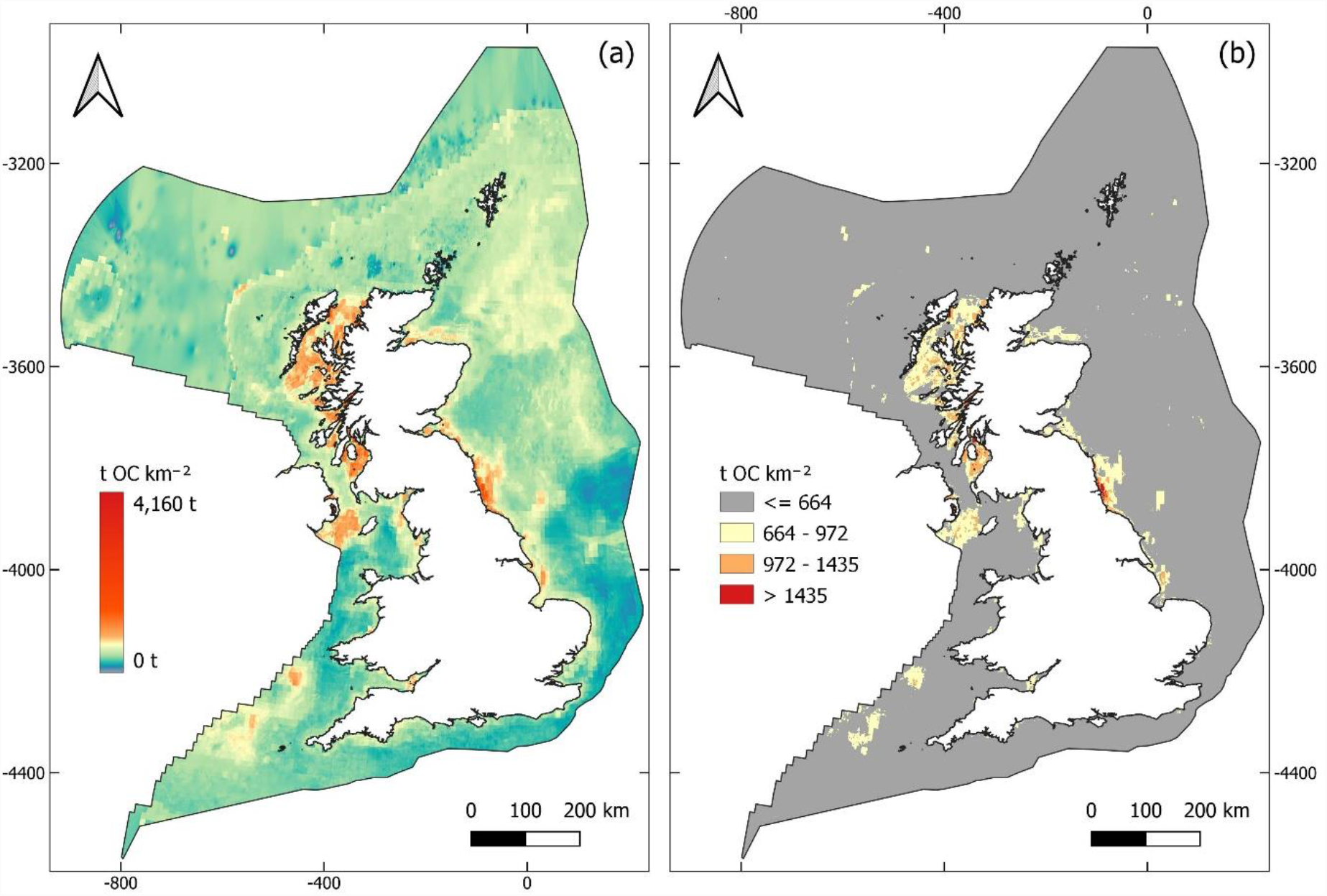
Seabed organic carbon (OC) stocks in the top 10 cm of sediment across the UK EEZ. **(A)** Concentration of OC across the study area. Colours follow an exponential scale from low concentrations in blue to high concentrations in red. **(B)** Areas of the study region which contain the highest concentrations of OC. Colours are equivalent to highest 0.1% of values - red, 1% - orange, and 5% - yellow.

Highest concentrations of 1435 - 4159 t OC km^-2^ (highest 0.1%) were predicted in the fjords and lochs of Scotland and Northern Ireland, and nearshore areas of Tyne and Wear (Northeast England) (Fig. 1b). High concentrations of 663 – 1433 t km^-2^ (highest 5% of values) were predominantly found in coastal and inshore waters, particularly around Scotland, Northeast England and the North Channel of the Irish Sea (Fig. 1b). Offshore areas with high concentrations of organic carbon were less extensive, predicted to occur in small areas to the East of the Bristol Channel, and on the shelf slope and in the North Sea (Fig. 1b). Together, these areas contain 10.1% of the total stock of organic carbon and occupy only 5.0% of the study region (Fig. 1b, Fig. 2a, Table 1). The remainder of the UK EEZ had lower organic carbon concentrations and less variation, ranging from ∼150-660 t km^-2^ and a mean of 382 ± 132 (SD) t km^-2^ (Fig. 1a).

**Table 1.**
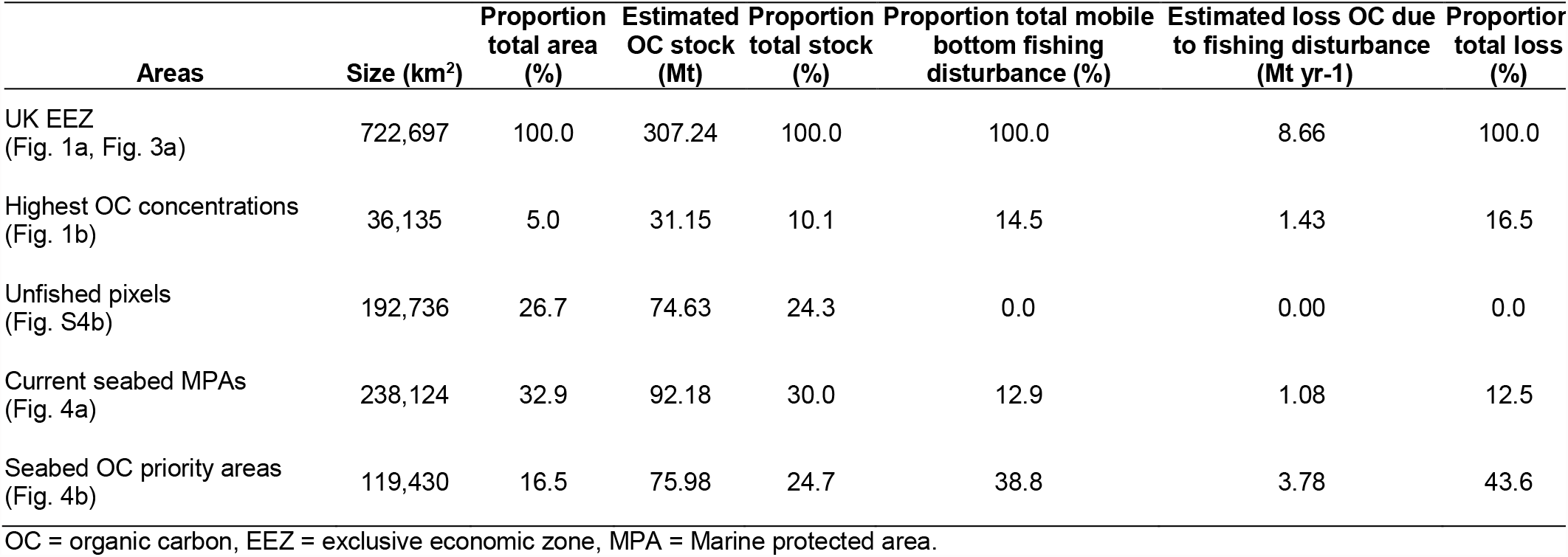
Summary statistics for different proportions of the study region. Information of the size, organic carbon (OC) stock, mobile bottom fishing disturbance, and potential losses of OC due to fishing disturbance in different areas of the UK EEZ.

| Areas | Size (km<sup>2</sup>) | Proportion total area (%) | Estimated OC stock (Mt) | Proportion total stock (%) | Proportion total mobile bottom fishing disturbance (%) | Estimated loss OC due to fishing disturbance (Mt yr<sup>-1</sup>) | Proportion total loss (%) |
| --- | --- | --- | --- | --- | --- | --- | --- |
| UK EEZ (Fig. 1a, Fig. 3a) | 722,697 | 100.0 | 307.24 | 100.0 | 100.0 | 8.66 | 100.0 |
| Highest OC concentrations (Fig. 1b) | 36,135 | 5.0 | 31.15 | 10.1 | 14.5 | 1.43 | 16.5 |
| Unfished pixels (Fig. S4b) | 192,736 | 26.7 | 74.63 | 24.3 | 0.0 | 0.00 | 0.0 |
| Current seabed MPAs (Fig. 4a) | 238,124 | 32.9 | 92.18 | 30.0 | 12.9 | 1.08 | 12.5 |
| Seabed OC priority areas (Fig. 4b) | 119,430 | 16.5 | 75.98 | 24.7 | 38.8 | 3.78 | 43.6 |

OC = organic carbon, EEZ = exclusive economic zone, MPA = Marine protected area.

**Fig. 2.**
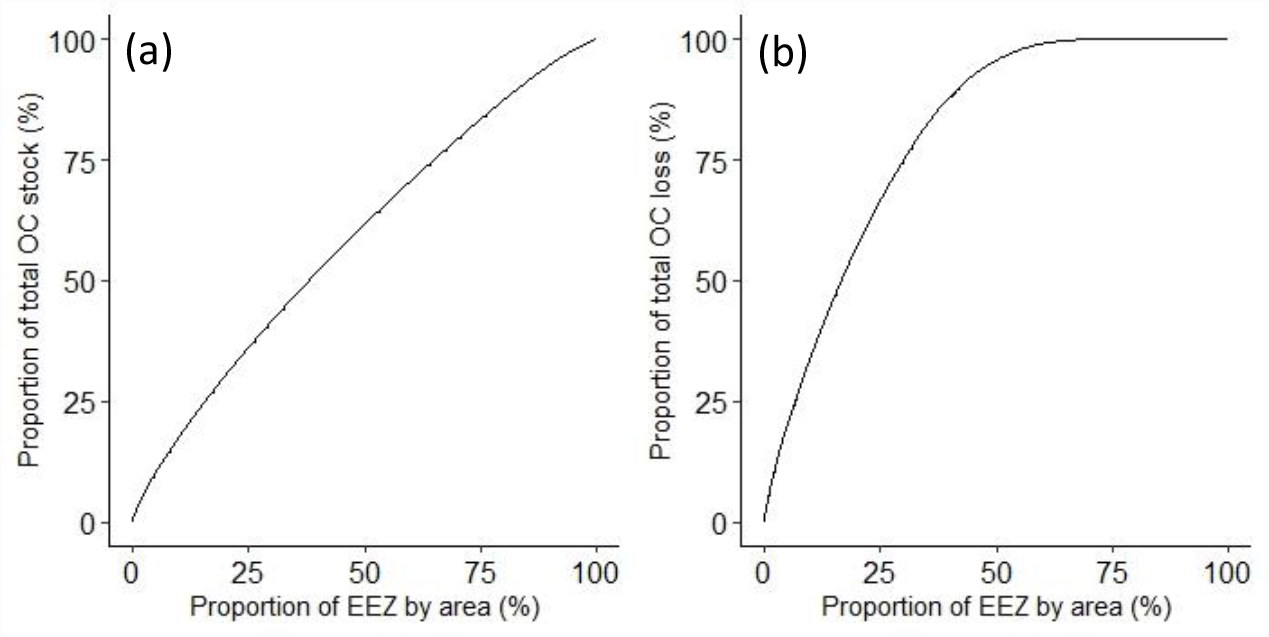
Cumulative distribution curves of organic carbon (OC) across the UK EEZ. **(A)** Cumulative distribution curve of estimated OC stock as a proportion of total stock across the study region. **(B)** Cumulative distribution curve of predicted OC loss due to mobile bottom fishing as a proportion of total OC loss across the study region.

### 2.2 Mobile bottom fishing disturbance

To identify where organic carbon is most at risk from mobile bottom fishing, fishing disturbance was quantified by combing spatial datasets of swept area ratios (SAR). Considering vessels with vessel monitoring systems (VMS) only (*37, 38*), i.e. > 15m long, the mean annual SAR from 2009 - 2018 ranged from <0.001 – 75.2 in fished pixels, with an overall mean SAR of 1.3 ± 2.5 across the study area (Fig. S1). Using spatial survey data of Scottish fishers with <15 m vessels (*39*) and data on fishing vessel footprints (*40*), a proxy SAR value was calculated for Scottish vessels lacking VMS. This ranged from 0.008 if 1 vessel was identified to fish in a given pixel, up to a SAR of 4.9 where 55 vessels stated they fished in that location. This added an additional 4.1% to the total fishing disturbance and 0.2% to the fished area calculated from VMS data alone (Fig. S2). Additionally, across 12 nautical miles (nm) in England and Wales, a proxy SAR value for smaller, inshore fishing vessels lacking VMS was estimated using sightings per unit effort data (SPUE) (*41*) and UK Fisheries Statistics (*42*). This led to a SAR ranging from 0.06 where SPUE was 0.001 up to a SAR of 61.6 where SPUE was 1.3. Overall, this added 8.3% to the estimated disturbance and 0.8% to the fished area as estimated by VMS data alone (Fig. S3) Combing these three datasets together, mean SAR across the study area was 1.4 ± 2.7, and ranged between <0.001 – 75.2 in fished pixels (Fig. S4a). A number of highly fished areas across the UK continental shelf and slope had SARs >5, while the most highly impacted area was in the English Channel, west of the Bassurelle Sandbank, with SARs >40 (Fig. S4a). Mobile bottom fishing occurred across the majority of the study area, with only 26.7% of the seabed estimated to be unaffected over the ∼10 year period covered by the dataset (Fig. S4b). Areas containing high organic carbon stocks were disproportionately affected by mobile demersal fishing, covering 5% of the study region but containing 14.5% of total fishing disturbance (Table 1, Fig. 1b).

### 2.3 Impact of fishing on organic carbon

To derive predicted values of organic carbon loss across the study region, estimates of organic carbon stocks and fishing disturbance were combined with published modelled values on the impact of mobile bottom fishing on organic carbon concentrations (*12*). The total potential loss of organic carbon from the top 10 cm of seabed was estimated as 129.9 Mt over 15 years, or 8.7 Mt of organic carbon per year (Fig. 3a, Table 1). Estimated potential losses per pixel ranged from 0 to 170 t OC km^-2^ yr^-1^, with highest losses predicted in similar areas with high organic carbon stocks (Fig. 1b, Fig. 3b). Further areas with high predicted loss were in the Fladen Grounds and East of Shetland, as well across much of the shelf slope and in coastal parts of East and South England (Fig. 3b). When selecting for the top 10% of estimated rates of loss, 34.0% of organic carbon loss were predicted from only 10.0% of the study region (Fig. 2b, Fig. 3b). The remainder of fished pixels had relatively low predicted organic carbon loss and less variation with 13.0 ± 9.5 t OC km^-2^ yr^-1^ (Fig. 3a).

**Fig. 3.**
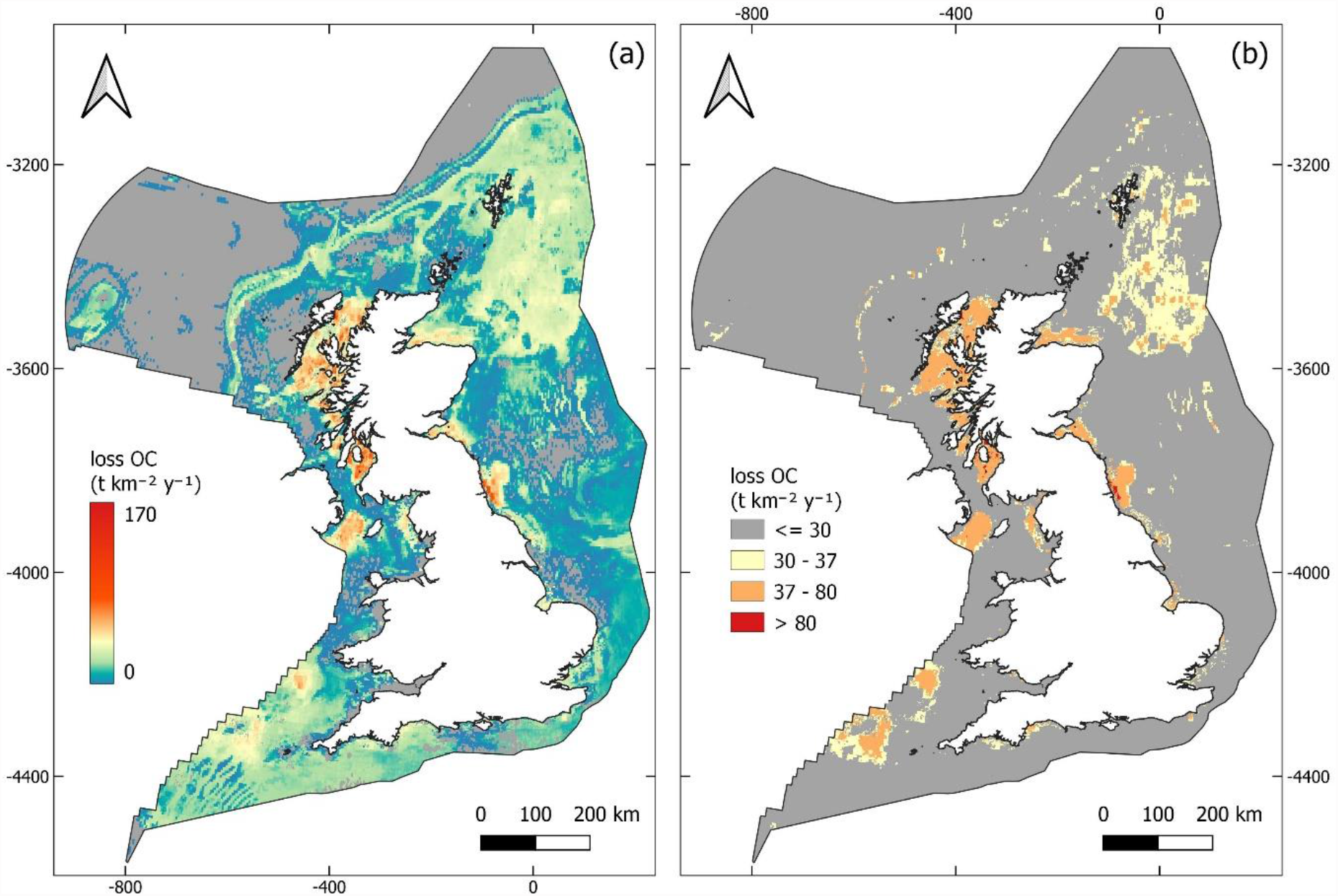
Predicted loss of seabed organic carbon (OC) stocks in the top 10 cm of sediment due to mobile bottom fishing across the UK EEZ. **(A)** Rates of OC loss across the study area. Colours follow a scale from low losses in blue to high losses in red. **(B)** Areas of the study area predicted to have the greatest rates of OC loss. Colours are equivalent to highest 0.1% of values - red, 5% - orange, and 10% - yellow.

### 2.4 Climate mitigation potential of the current seabed MPA network

At the time of writing there were 231 MPAs designated in the UK which included seabed components (Fig. 4a). These include sites designated as Special Areas of Conservation (SAC), Marine Conservation Zones (MCZ) and Nature Conservation Marine Protected Areas (NCMPA). In total these MPAs cover 32.9% of the study region, 30.0% of total organic carbon stock, 12.9% of fishing disturbance, and 12.5% of the predicted organic carbon loss (Table 1). Even though 32.9% of the study region is under a protected area designation, <1% is fully or highly protected from fishing (*20*), eliminating the use of mobile fishing gears. If the entire current MPA network were to be closed to mobile bottom fishing, there would be a predicted net saving of 0.59 Mt OC yr^-1^ or 6.9% of the estimated loss due to current fishing disturbance (after accounting for mean modelled fisheries displacement) (Fig. 4a).

**Fig. 4.**
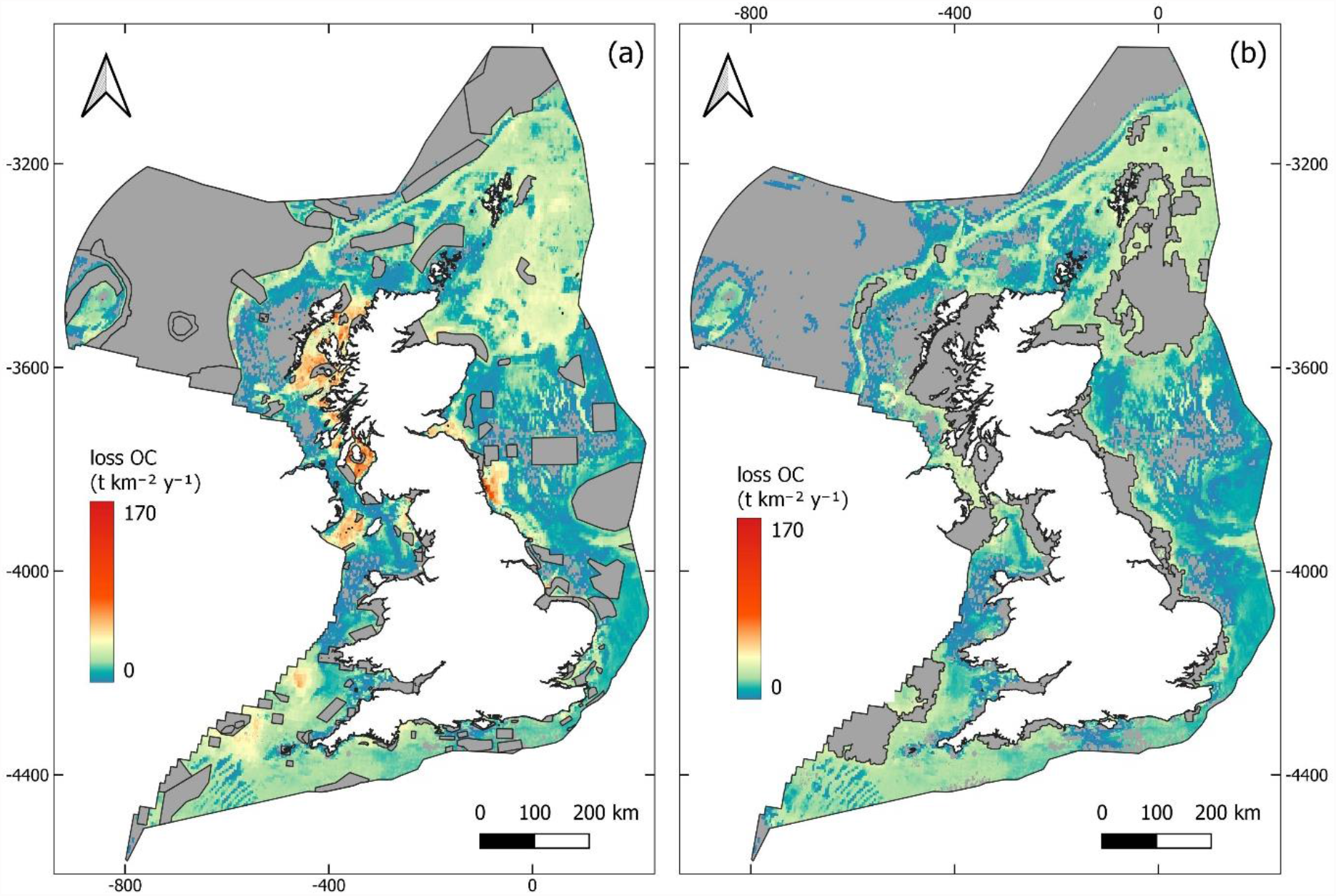
Predicted loss of organic carbon (OC) following mean modelled fisheries displacement if given areas were closed to mobile bottom fishing. **(A)** Predicted loss of OC if mobile bottom fishing was excluded from all currently designated seabed MPAs. **(B)** Predicted loss of OC if all seabed OC priority areas were closed to mobile bottom fishing. Colours follow a scale from low concentrations of OC loss in blue to high concentrations in red.

### 2.5 Priority areas to protect seabed organic carbon for climate mitigation

To identify where protection from mobile bottom fishing would achieve the largest savings of organic carbon, areas were selected if they contained aggregations of fished pixels with high concentrations of predicted organic carbon loss or organic carbon stock (Fig. S5, Fig. S6, Table S1). These were refined to a final set of 33 priority areas after accounting for the estimated net organic carbon saving in each area following mean modelled fisheries displacement (Fig. 4b, Fig. 5). In total the priority areas cover 16.5% of the study region, 24.7% of total organic carbon stock and 43.6% of the predicted organic carbon loss due to mobile bottom fishing (Table 1, Fig. 4b, Table S2). After accounting for mean fisheries displacement (Fig. S7, Fig. S8) the predicted net saving of organic carbon from establishing mobile bottom gear closures across all priority areas was 2.56 Mt OC yr^-1^ or 29.6% of the estimated organic carbon loss across the study region (Fig. 4b).

**Fig. 5.**
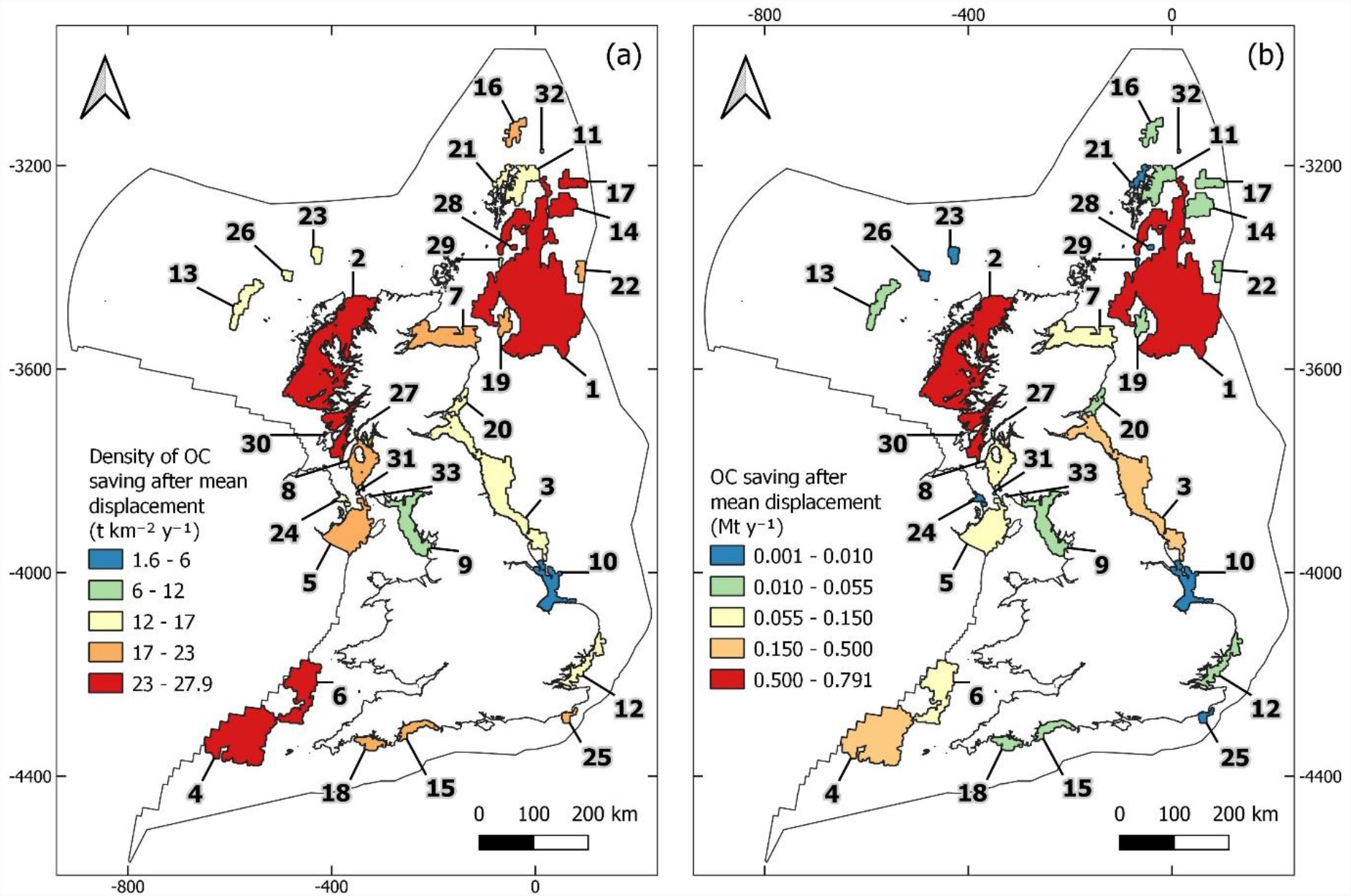
Estimated net organic carbon (OC) saving if each seabed OC priority area were to be closed to mobile bottom fishing. **(A)** The density of net organic carbon (OC) saving, and **(B)** the total net OC saving estimated following mean fisheries displacement. Numbered labels match “Proposed priority area” IDs shown in Table S2.

Based on highest densities or total net savings of organic carbon, ten priority areas dominated the potential network benefit (Fig. 5, Table S2). In total these ten areas (#1-8, 14, 27) make up 87.2% of the network benefit, predicted to save 25.8% of total organic carbon loss across the study region and cover only 13.2% of the area (Fig. 5, Table S2). The remaining 23 priority areas save comparatively little organic carbon, on average 0.2 ± 0.1% of total organic carbon loss, cumulatively saving 3.8% of total loss across the region, while covering 3.3% of the area (Fig. 5b, Table S2).

## 3. Discussion

The marine environment represents a large fraction of the UK organic carbon store. Seabed sediments across the UK EEZ are estimated to store 307 Mt of organic carbon in their top 10 cm (Fig. 1a), which compares with an estimated 550 Mt stored in high carbon terrestrial and intertidal vegetation and soils to a depth of 30 cm (*43*). Many of these habitats, such as peatland, forests, marsh, mudflats and heathland, are being recognised as important carbon stores with the potential to mitigate against climate change, which has led to widespread efforts to protect and restore them globally (*44*). The contribution of seabed sediment habitats, by contrast, is under-appreciated, but due to the scale of their organic carbon stores, there is need to urgently reconsider their management and protection.

Across the UK EEZ the potential for loss of organic carbon from seabed sediments due to disturbance from mobile bottom fishing was found to be high. Total loss from the top 10 cm of seabed was estimated as 8.7 Mt of organic carbon per year (Fig. 3a). If this was entirely remineralised it would equal 32 Mt of CO_2_, equal to ∼9% of UK CO_2_ emissions (*45*). Although some of this organic carbon is likely to be remineralised to CO_2_, some will simply be transported, either being consumed or relocated elsewhere (*11, 46*). Even if organic carbon is remineralised to CO_2_ it does not mean that it will be released to the atmosphere or even stay in the form of aqueous inorganic carbon within the ocean; it is expected that a proportion would be re-fixed through photosynthesis of marine algae and aquatic plants (*18*). Additionally, organic carbon in the top 10 cm of the seabed is still undergoing active processing with natural remineralisation occurring across differing sediment depths dependent on environmental settings (*47*). Therefore, the total mass of organic carbon stored over climate relevant timescales is expected to be less than the total present. The oceanic carbon cycle is highly complex (*6*), how depletion of seabed sediment organic carbon will effect marine carbon cycling and the atmospheric concentration of CO_2_ requires further research (*18*). Even so, the protection and restoration of organic carbon stores across land and sea is widely accepted as necessary for climate change mitigation (*44*).

The current UK seabed MPA network was designed for the protection and restoration of biodiversity while minimising economic impact and disruption to fisheries activities (*48*). Seabed sediment carbon was not considered, and in this study we have shown that the goal of carbon protection is poorly served by the present MPA configuration. The current network of 231 seabed MPAs covers 32.9% of the study region but only 30.0% of total organic carbon stock, 12.9% of fishing disturbance, and just 12.5% of the predicted organic carbon loss (Fig. 4a, Table 1). If the entire network were to be closed to mobile bottom fishing, with mean modelled fisheries displacement, there would be a predicted net saving of only 0.59 Mt organic carbon yr^-1^ or 6.9% of the estimated total loss due to current fishing disturbance (Fig. 4a).

Further expansion of the MPA network seems unlikely in the near future, given that it is already among the most extensive in the world at 38% total coverage (*32*). However, other legislative instruments for area-based management of mobile bottom fishing could be used to reduce impact on seabed sediment organic carbon stores. In the UK, fisheries by-laws enacted through government agencies could close specific areas to mobile bottom fishing while allowing less damaging fishing activities to continue, without the need to expand the current MPA network. These simpler regulatory tools may allow local agencies or devolved governments to more quickly protect organic carbon stores.

The priority areas identified in this study could act as sites for further investigation and potentially new management measures for the protection of seabed organic carbon. They cover 24.0% of the total organic carbon stock, 43.0% of the predicted organic carbon loss and only 16.0% of the UK EEZ (Fig. 4b, Table 1). If mobile bottom fishing was entirely excluded from priority areas, mean modelled fisheries displacement would predict a net saving of 2.56 Mt OC yr^-1^ or 29.5% of the estimated total loss due to current fishing disturbance (Fig. 4b).

The majority of the organic carbon saving from priority areas was predicted to come from the establishment of three large closed areas (Table S2). The first covered the Fladen Grounds and surrounding areas (#1), the second covered the inland sea lochs, inlets and waterbodies West of Scotland (#2), and the third around Haig Fras (#4) (Fig. 5b). Together these three areas save 18.6% of the total organic carbon loss across the study site or 62.9% of the total estimated benefit of closing all 31 priority areas, and cover only 8.8% of the UK EEZ (Table S2). It is also important to consider areas with high densities of organic carbon saving as they would allow for the largest climate mitigation benefit in the smallest possible cumulative area. The priority areas with the highest density of organic carbon savings following mean modelled fisheries displacement were Loch Fyne, Haig Fras, West of Scotland, North Fladen 1, and Celtic Deeps, (# 27, 4, 2, 14 6) (Fig. 5a, Table S2). Together these five areas save 11.7% of the total organic carbon loss and cover only 5.2% of the UK EEZ (Table S2).

Protecting areas of seabed sediment would appear to be an important climate mitigation strategy, however carbon has not been considered in national management plans. As seen in this study, areas of seabed sediments with high organic carbon concentrations are often disproportionately impacted by fishing disturbance due to the presence of highly productive fishing grounds. In the UK EEZ sediments with high carbon concentrations cover only 5% of the region, but contain 14.5% of total mobile demersal fishing disturbance (Table 1, Fig. 1b). To gain significant climate mitigation benefit, fishing activities within these areas will need to transition to less disruptive methods. Where national MPA networks have already been established, carbon considerations will need to be retrofitted to current management schemes, and further protections are likely to be necessary outside of MPA networks. Countries that are still developing their marine management plans should consider seabed sediment organic carbon as part of their MPA development processes.

Our estimated loss of 8.7 Mt of organic carbon per year due to mobile bottom fishing is in accord with a global study which estimated UK losses as 5.2-12.8 Mt per year (*11*). However, there remains significant uncertainty in the magnitude and direction of mobile fishing effects (*8, 13, 19*). Our estimates rely on one modelling exercise undertaken in the North Sea which was parametrised by in-situ data from five locations but the impact of mobile bottom fishing was only simulated (*12*). Their model outputs predict that in all settings mobile bottom fishing would reduce organic carbon concentrations in seabed sediments (*12*). However, in areas with high hydrodynamic activity, low deposition rates, large oxygen penetration depths and highly refractory organic carbon, the effect of fishing disturbance may be limited (*13*). Our calculated losses of organic carbon therefore come with significant uncertainty. For the spatial prioritisation exercise however, higher confidence can be assumed, with areas containing high organic carbon concentrations and high fishing pressure generally expected to have highest loss of organic carbon even if the magnitude is yet to be fully resolved.

As the mean fisheries displacement scenario estimates that only 29.6% of losses would be saved by closing all priority areas to mobile bottom fishing we must also consider whether additional non-spatial management measures may be necessary (*29, 49*). As the concentration of seabed sediment organic is relatively similar across much of the study site (Fig. 1a, Fig. 2a), any management measures which significantly reduce overall seabed disturbance would be expected to have carbon benefit. For example if changes were made across the mobile bottom fishing fleet with gear modifications to reduce seabed disturbance, overall effort reductions through total effort quotas, and the incentivisation to switch to alternative fishing methods (*49*), the potential organic carbon savings could be significant.

Although not a focus of this study, other activities which also impact the seabed, including marine energy developments, mineral extraction, shipping and coastal development should also be considered for their potential to impact seabed sediment organic carbon stores (*8, 16*). The scale of impact is expected to be highly site specific and activity dependent, but any human activity which increases the distribution or frequency of seabed sediment disturbance may limit surface organic carbon concentrations and subsequent organic carbon storage (*8, 46*). Even so, mobile bottom fishing is by far the most widespread human activity affecting the seabed and therefore likely dominates cumulative human impacts on marine organic carbon stores (*14, 15*).

There is an urgent need to decarbonise our industries, requiring a step change in how we extract and use resources from our planet (*50*). We must concurrently carbonise our environment by protecting organic carbon stores and promoting natural carbon sequestration on both land and at sea (*28*). The UK fishing industry is estimated to emit ∼914.4 kt of CO_2_ per year (*51*), but its potential emissions from disturbance to sediment organic carbon stores could greatly outweigh these direct emissions. Our study shows that targeted area-based management of mobile bottom fishing has the potential to provide significant climate mitigation benefit by protecting organic carbon in seabed sediments. Due to previous management priorities, much of this climate mitigation potential is likely to occur outside of current MPA networks. We must re-evaluate current seabed management plans and incorporate new evidence-based carbon considerations.

## 4. Materials and Methods

### 4.1 Analysis software

All analyses were undertaken in R 4.0.5 (*52*). Manipulation and display of raster and spatial vector data was carried out using the *raster* and *sf* packages respectively (*53, 54*). The *fasterize* package was used to convert spatial polygon data into raster layers for further manipulation (*55*) and the *ggplot2* package was used for plotting graphical data (*56*). Further editing and display of spatial data was also carried out in QGIS (*57*). A list of input data sources required to run the analyses is shown in Table S4.

### 4.2 Seabed sediment organic carbon distribution and standing stock

Due to data availability, estimates of organic carbon standing stock in seabed sediments across the UK EEZ were constrained to the top 10 cm of the seabed. Four previously published spatial analyses of organic carbon density were identified, each covering varying spatial extents and resolutions across the Northeast Atlantic (*33-36*). All datasets, except Diesing, Kroger, Parker, Jenkins, Mason and Weston (*33*), were available as estimated organic carbon density (kg m^-2^) in each spatial pixel for their respective extents. Publicly available data outputs from Diesing, Kroger, Parker, Jenkins, Mason and Weston (*33*) only contain estimates for the concentration of organic carbon as a percent of sediment dry weight. This was converted to density values (kg OC m^-2^) using data from Smeaton, Hunt, Turrell and Austin (*35*) who modelled dry bulk density (kg dry sediment per m^3^) across the study area. Final estimates of organic carbon in kg m^-2^ were made at a resolution of 500 × 500 m, as this is approximately the highest resolution of the 4 published datasets. All datasets were projected onto a 500 m resolution grid covering the UK EEZ with Lambert Azimuthal Equal Area coordinate system. Projections used bilinear interpolation and the mean concentration of organic carbon across the datasets was calculated where overlap occurred. Concentration data (kg OC m^-2^) was converted to t km^-2^ for further analysis.

### 4.3 Mobile bottom fishing disturbance

Data on the extent and disturbance from mobile bottom fishing was primarily derived from published outputs by the International Council for the Exploration of the Sea (ICES). ICES calculate the annual surface swept area from mobile bottom fishing as hours fished within a given year multiplied by average fishing speed multiplied by gear footprint (*38*). The annual swept area ratio (SAR) is the sum of the swept areas divided by the area of each grid cell (*38*). In ICES (2019)(*38*) the total SAR from mobile bottom fishing vessels (otter trawls, bottom trawls, dredges and seines) carrying vessel monitoring systems (VMS) registered to European vessels (Belgium, Denmark, France, Germany, Ireland, The Netherlands, Norway, Sweden and the UK) was available for each year from 2009 to 2017 at a resolution of 0.05° (*38*). The mean SAR across years was calculated to gain a mean annual SAR for each pixel.

Where data from the ICES (2019)(*38*) publication were lacking for parts of the study area, a less precise average annual SAR value was derived from ICES (2021)(*37*). Here, published values of mean SAR per year were available at a resolution of 0.05° for the period 2013-2018 from VMS data provided by all EU vessels, United Kingdom, Faroes, Iceland, and Norway (*37*). As SAR data in ICES(2021)(*37*) were only available as discrete categorical values, the median value within each categorical range was used where ICES (2019)(*38*) data were lacking.

As ICES data only considers those vessels with VMS, a large portion of the inshore, <15m fishing fleet, will not be considered in the calculation of SAR. A conservative proxy SAR value for <15 m fleet was calculated using survey data from ScotMap in Scotland (*39*), and CEFAS data in England and Wales (*41*). Using surveys of fishers who own <15 m Scottish registered vessels, ScotMap contains spatial data displayed at 0.025° resolution, on the number of vessels which identified that they carry out bottom mobile fishing activities within different areas across Scotland’s marine zone. Integer values for the number vessels are displayed for three vessel categories (*Nephrops* trawls: trawler vessels primarily targeting *Nephrops norvegicus*; Other trawls: any other trawler vessel; and Dredges: any type of bottom dredge). Where pixels contain <3 vessels, data are only shown as a categorical value (1-3 vessels), therefore a conservative value of one vessel was used here for further analysis. Number of vessels in Scotland’s marine area only were converted to estimated SARs using data from Eigaard, Bastardie, Breen, Dinesen, Hintzen, Laffargue, Mortensen, Nielsen, Nilsson, O’Neill, Polet, Reid, Sala, Sköld, Smith, Sørensen, Tully, Zengin and Rijnsdorp (*40*). Average values of hourly swept area (SA: km^2^ h^-1^) were estimated as 1.2, 0.5 and 0.1 for Nephrops trawls, Other trawls and Dredges respectively (*40*). Using a conservative estimate that each vessel passes through each pixel once per year, and that vessels are assumed to travel at approximately 3 knots while fishing (*40*) (and are therefore estimated to travel through 2.75 pixels per hour), the number of vessels per pixel was converted to an average annual SAR using the formula: SAR = n*(SA/2.75)/A; where n is the number of vessels and A is the estimated surface area of the pixel calculated using the *area* function from the *raster* package.

To estimate inshore fishing effort in English and Welsh waters, Vanstaen and Breen (*41*) calculate a value of sightings per unit effort (SPUE) at a resolution of 0.05° x 0.025° using CEFAS sightings data from fisheries monitoring vessels and overflight in 2007-2009 and 2010-2012. The value of SPUE for all vessels with mobile gears within 12 nm was used here as a proxy for effort from the <15 m bottom mobile inshore fleet. Although these data may contain some vessels with VMS, and some mobile pelagic vessels, in England and Wales the <15 m fleet makes up over 90% of registered fishing vessels, and activity days data for vessels under 15 m length shows that just 1.76% of all fishing activity days for trawlers was undertaken using midwater trawls, and so 98.24% of activity days by trawlers is expected to be using bottom gears (*41*). A mean of the two datasets (2007-2009 & 2010-2012) was taken, and mean SPUE was converted to a coarse value of estimated annual SAR by rescaling the data so the total SAR in English and Welsh waters was equal to 1.73 times the total SAR calculated above for Scottish <15 m vessels. The 1.73 value was derived from the MMO 2019 UK Fisheries Statistics as the mean difference in total capacity of English and Welsh small vessels compared to Scottish vessels in 2016-2019 (*42*).

All SAR values were projected to align with organic carbon stock data using bilinear interpolation. As SAR values do not account for sub-pixel variation in fishing patterns this assumes that SAR is spread evenly across all 500m pixels that lie within each SAR pixel. A final total SAR value per pixel was calculated by summing the combined ICES data layer with ScotMap and CEFAS SAR estimates. Final SAR values were used to indicate the theoretical number of times the entire pixel has been impacted by mobile bottom fishing if effort was evenly distributed within that pixel (*37, 38*). Therefore, as an example, a SAR of 3 was assumed to mean that the entire pixel is trawled three times per year, while a SAR of 0.4 was assumed to mean that 40% of the pixel is trawled once per year.

### 4.4 Impact of fishing on organic carbon

The impact of mobile bottom fishing on seabed sediment organic carbon stocks was taken from De Borger, Tiano, Braeckman, Rijnsdorp and Soetaert (*12*). This study models the estimated percentage change in organic carbon concentration in the top 10 cm of sediment from repeated fishing events by a tickler-chain beam trawl over 15 years within the North Sea. Although this is an estimate for only one gear type, the penetration depth was modelled as 3.2 (1.5-6.7) cm and 1.9 (1.0-3.7) cm for mud and sand respectively; which is around the average disturbance expected for the mobile bottom fishing fleet in this study (*40*).

De Borger, Tiano, Braeckman, Rijnsdorp and Soetaert (*12*) models predicted impact over three sediment types (Coarse sediments, Fine sands and Muds) and two levels of nutrient concentration (High and Low) – leading to five different habitat types (Coarse, FineL, FineH, MudL and MudH). To align with De Borger, Tiano, Braeckman, Rijnsdorp and Soetaert (*12*), each pixel of the study area was classified to these five scenarios based on their predicted sediment type (Folk category) from Smeaton, Hunt, Turrell and Austin (*35*) and distance from shore (Table S3, Fig. S9). Those pixels predicted to be rock and boulders were assumed to have no fishing impact on carbon (Table S3, Fig. S9).

The outputs from De Borger, Tiano, Braeckman, Rijnsdorp and Soetaert (*12*) used in this study are the values of modelled percent loss in organic carbon over 15 years due to mobile bottom fishing at intensities of 1,2,3,4 and 5 trawl passes per year for each habitat respectively. In each pixel in the study area, the SAR values calculated above were rounded to the nearest integer to match De Borger, Tiano, Braeckman, Rijnsdorp and Soetaert (*12*) scenarios and the estimated loss of organic carbon in each pixel was calculated by multiplying the organic carbon stock estimates by their respective modelled percent loss value (Table S5). For pixels with SAR < 1, the organic carbon stock value was first multiplied by the SAR value (so only a proportion of the pixel was being considered) and was then treated the same as a SAR value of 1 (meaning a proportion of the pixel is considered to be fished once per year) (Table S5). As De Borger, Tiano, Braeckman, Rijnsdorp and Soetaert (*12*) did not produce modelled outputs for scenarios of over 5 trawl passes per year, pixels with SAR > 5 were treated the same as pixels with SAR >= 4.5. This may lead to an underestimate of impact, however the level of impact from each additional trawl pass is expected to reduce at higher intensities (*11, 12*). As De Borger, Tiano, Braeckman, Rijnsdorp and Soetaert (*12*) estimate the loss of organic carbon over 15 years, a final value of organic carbon lost per year was calculated for each pixel by dividing the total loss of organic carbon by 15. Although this has the underlying assumption that organic carbon is lost uniformly each year, the true trajectory of loss is unknown and therefore this measure of average loss per year is an approximate metric.

### 4.5 Current seabed MPA network

Data on the distribution and classification of designated seabed MPAs were gathered from individual government agencies. This included Special Areas of Conservation in all UK waters (*58*), Marine Conservation Zones in England (*59*), Northern Ireland (*60*) and Wales (*61*), as well as Nature Conservation MPAs in Scotland (*62*). All MPAs were collated and any MPAs whose protected features contained no biological seabed components were excluded from further analysis. MPAs were excluded if they only protected pelagic or mobile species (e.g. birds, cetaceans, seals, pelagic fish), and/or geomorphological or oceanographic features (e.g. fronts, quaternary of Scotland). These data were used to assess the overlap of the current network with organic carbon stocks and losses. Additionally, the predicted net effect on organic carbon from establishing mobile bottom gear closures across all MPAs was calculated using mean fisheries displacement as described below.

### 4.6 Impact of fisheries displacement

It is challenging to predict how fishers will adapt to new spatial management measures and therefore difficult to calculate the true impact of fisheries displacement. Some fishing effort will be eliminated as fishers move to alternative methods or industries (*30*). Effort that is displaced rather than eliminated is likely to move to areas both proportional to current effort, due to the distribution of productive fishing grounds, and inverse exponentially from the closed area, to minimise displaced distance and gain benefits from spill-over effects (*30, 31, 63, 64*). Maximum displaced distance is also likely to vary dependent on the size of the vessel and distance from available ports (*27, 63, 64*). It is for these reasons the mean effect of multiple simulated fisheries displacement scenarios was used in this study to best estimate net organic carbon savings (Fig. S7, Fig. S8).

To calculate the effect of excluding mobile bottom fishing from given areas, fisheries disturbance (SAR) was recalculated using five displacement scenarios. The first scenario assumed that closing an area to mobile bottom fishing would lead to complete elimination of fishing effort. In this case SAR was equal to zero in all pixels within the closed area and SAR would remain constant in all areas outside the closure. The remaining four scenarios were based on two displacement models and two maximum displacement distances. The two displacement models are: 1) fishing disturbance spread proportionally across a given area based on current fishing disturbance; 2) disturbance spread inverse exponentially across a given area with decreasing disturbance as distance from the closed area increases (see Hoos, Buckel, Boyd, Loeffler and Lee (*63*) for details). The first model assumes that the current distribution of fishing disturbance reflects the locations of productive and unproductive fishing grounds and is therefore likely to be targeted in similar levels by displaced disturbance. In this model, fisheries disturbance in the selected area outside the closed area would occur in the same ratios but at higher magnitudes than before the closed area was introduced (*31*). The second scenario is equivalent to “fishing the line”, whereby fishers are expected to minimise displaced distance while gaining potential benefit from spill-over effects from the closed area (*63, 64*). The two displacement models were calculated over maximum displacement distances of 50 km and 200 km surrounding each closed area, to model both local and regional displacement.

Except for in the elimination scenario, all displacement scenarios assume that 100% of the fishing disturbance is displaced to new locations within the study region but fishing cannot be displaced into other modelled closed areas. Also, it is assumed that fishing cannot be displaced into unfished pixels because the data would suggest that if these areas have been completely avoided by fishers for ∼10 years, they are likely to be inappropriate for mobile bottom fishing activities. It should also be noted that the SAR data calculated above were taken to represent areas which are open or closed to mobile bottom fishing. It is possible that some areas across the study region have seasonal or year-round closures to mobile bottom fishing but the SAR data indicates that they are fished. Other legislative instruments or area closures which limit mobile bottom fishing are not considered in this study.

Finally, the mean of the five scenarios was taken across the study region and the predicted loss of organic carbon was recalculated as above. The net effect of closure was calculated as the difference in organic carbon lost between current fishing patterns and the mean displacement scenario.

### 4.7 Selecting potential seabed organic carbon priority areas

Sites were selected as potential priority areas if they were predicted to have high current or future organic carbon loss and therefore removal or protection from mobile bottom fishing pressure would have most significant carbon savings. Pixels with the highest 10% of predicted organic carbon loss were combined with pixels with the highest 5% of organic carbon stock. Unfished pixels were removed from selection as these would show no direct additionality from protection. Spatial resolution of analysis was reduced to 2 km x 2 km pixels to reduce processing time and to give coarser area selections. Neighbouring pixels were joined using an 8-way Queen’s case method and raster data were then converted to individual polygons. A 2 km buffer was drawn around each polygon and overlapping polygons were dissolved to form single areas. Finally, any holes were removed from each area using the *remove*.*holes* command from the *spatialEco* package. The potential priority areas were refined by selecting only those which met one of the following four criteria: 1) contained at least 0.1% of total predicted organic carbon loss across the study region; 2) density of organic carbon loss within the top 10% of all selected areas; 3) contained at least 0.1% of total organic carbon stock across the study region; 4) density of organic carbon stock within the top 10% of all selected areas.

### 4.8 Proposed seabed organic carbon priority areas

Potential priority areas were further refined to a set of final proposed areas after accounting for mean modelled fisheries displacement. Each individual area was dropped in turn and the effect on organic carbon after mean fisheries displacement from all other areas was recalculated. Based on this “drop one” estimate, priority areas were eliminated if they contributed to a saving of less than 0.1% of the total organic carbon loss across the study region or their predicted loss rate was less than the mean of all individual potential priority areas. From this final set of proposed priority areas, the net effect of establishing mobile bottom gear closures across all areas was then recalculated using mean modelled fisheries displacement.

Finally, for each proposed priority area the net effect on organic carbon was calculated in two ways. Firstly, as conducted for the potential priority areas, their contribution was estimated using mean fisheries displacement and a “drop one” method. Secondly, mean fisheries displacement was calculated for a scenario where only the individual priority area is closed alone, and the net effect on organic carbon taken as the difference between this scenario and organic carbon lost with current fishing patterns. The mean of these two methods was taken as an estimated organic carbon benefit from closing each individual priority area.

## 6. Acknowledgments

## Funding

This work was funded by BLUE Marine Foundation through the Barclays Ocean Climate Impact grant.

## Author contributions

Conceptualization: GE

Methodology: GE

Investigation: GE

Visualisation: GE

Supervision: CMR

Writing—original draft: GE

Writing—review & editing: CMR, GE

## Competing interests

The authors declare that they have no conflict of interest.

## Data availability statement

All data and associated code will be made available in the supplementary material or in the Pangea data repository upon publication.

## Notes

### Competing Interest Statement

The authors have declared no competing interest.

