## Supplementary material for "Protecting seabed sediment carbon for climate mitigation: a UK case study": Fig. S

**This PDF file includes:**

Figs. S1 to S9

**Other Supplementary Materials for this manuscript include the following:**

Data Table S1 to S5

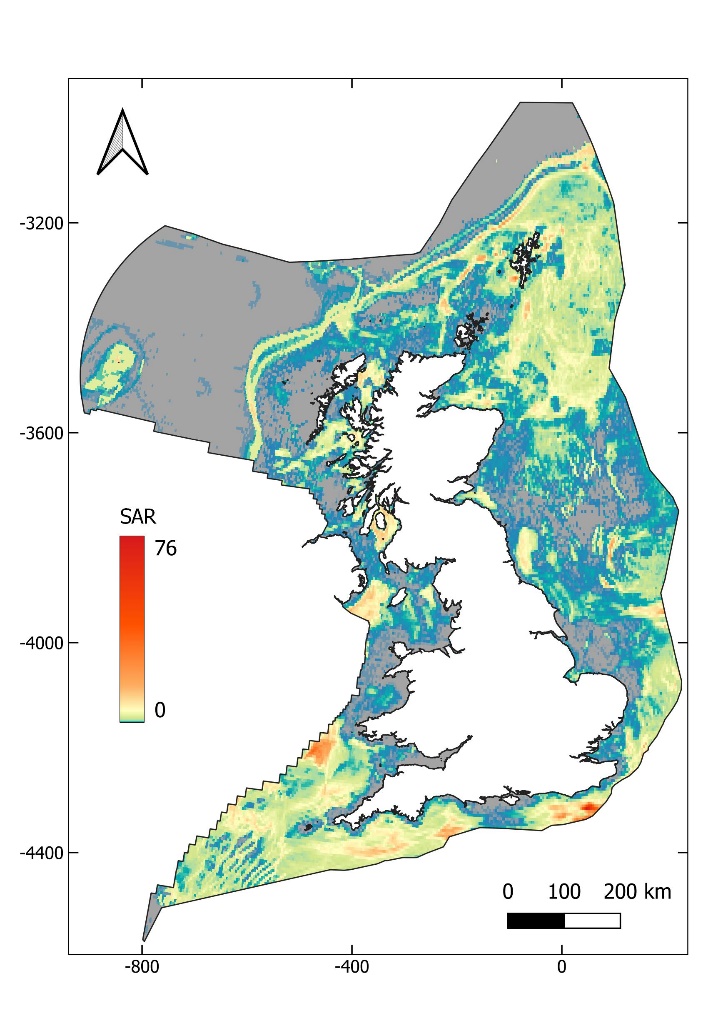

**Fig. S1** Swept area ratio from vessels with vessel monitoring systems (VMS) only (*1, 2*)

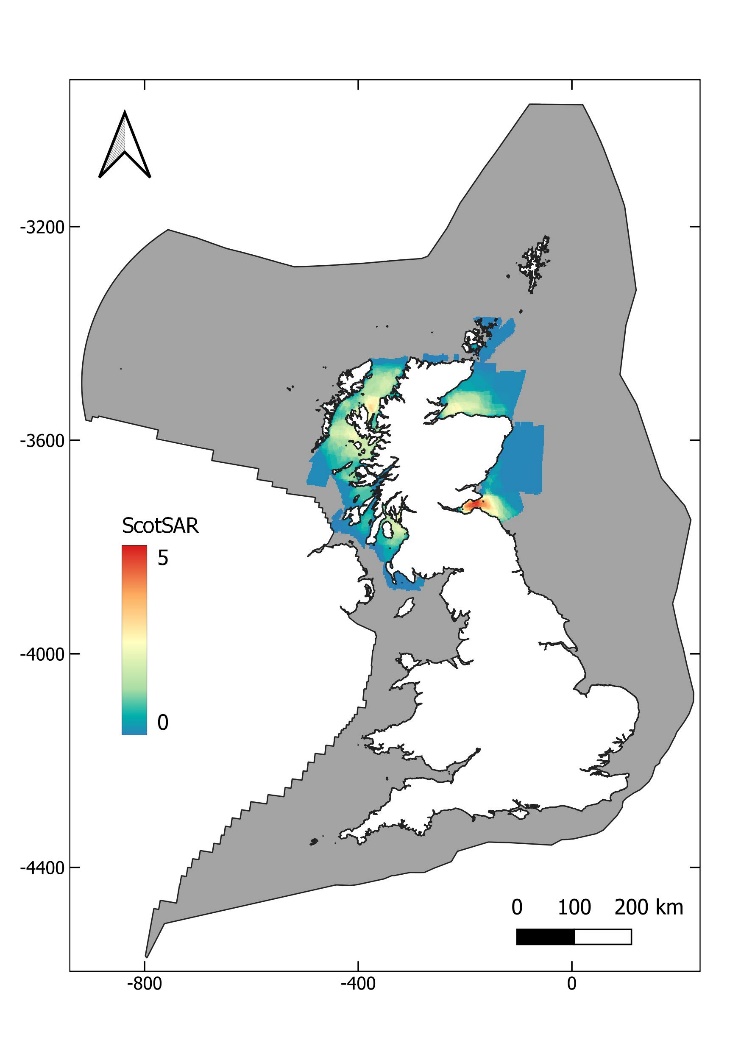

**Fig. S2** Swept area ratio for non-VMS (vessel monitoring system) vessels, calculated from ScotMap (*3*).

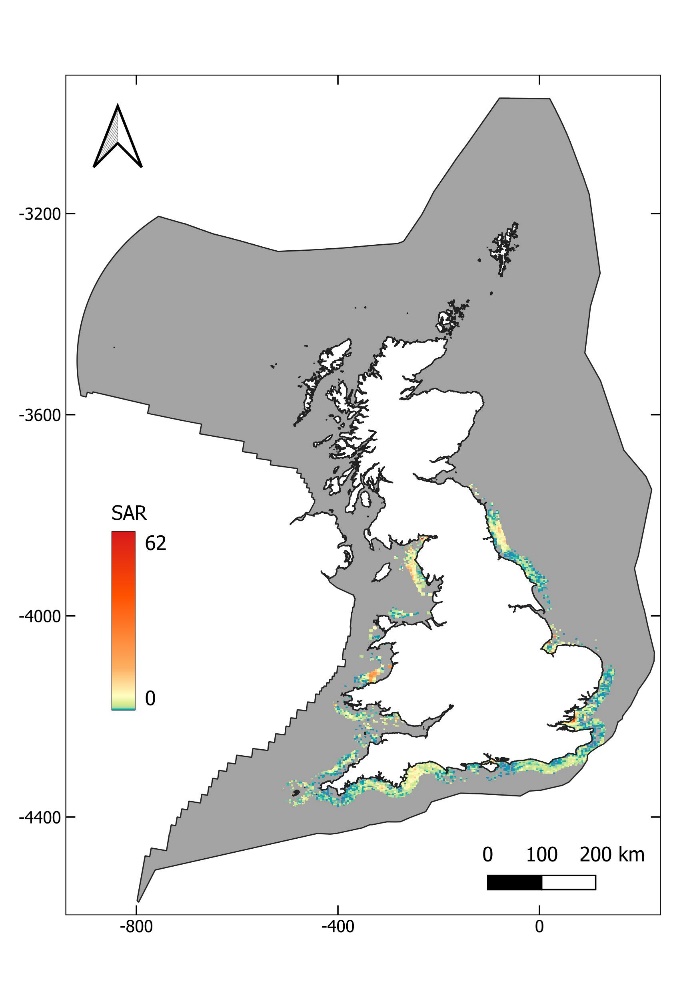

**Fig. S3** Swept area ratio calculated for non-VMS (vessel monitoring system) vessels, from CEFAS sightings per unit effort data (*4*).

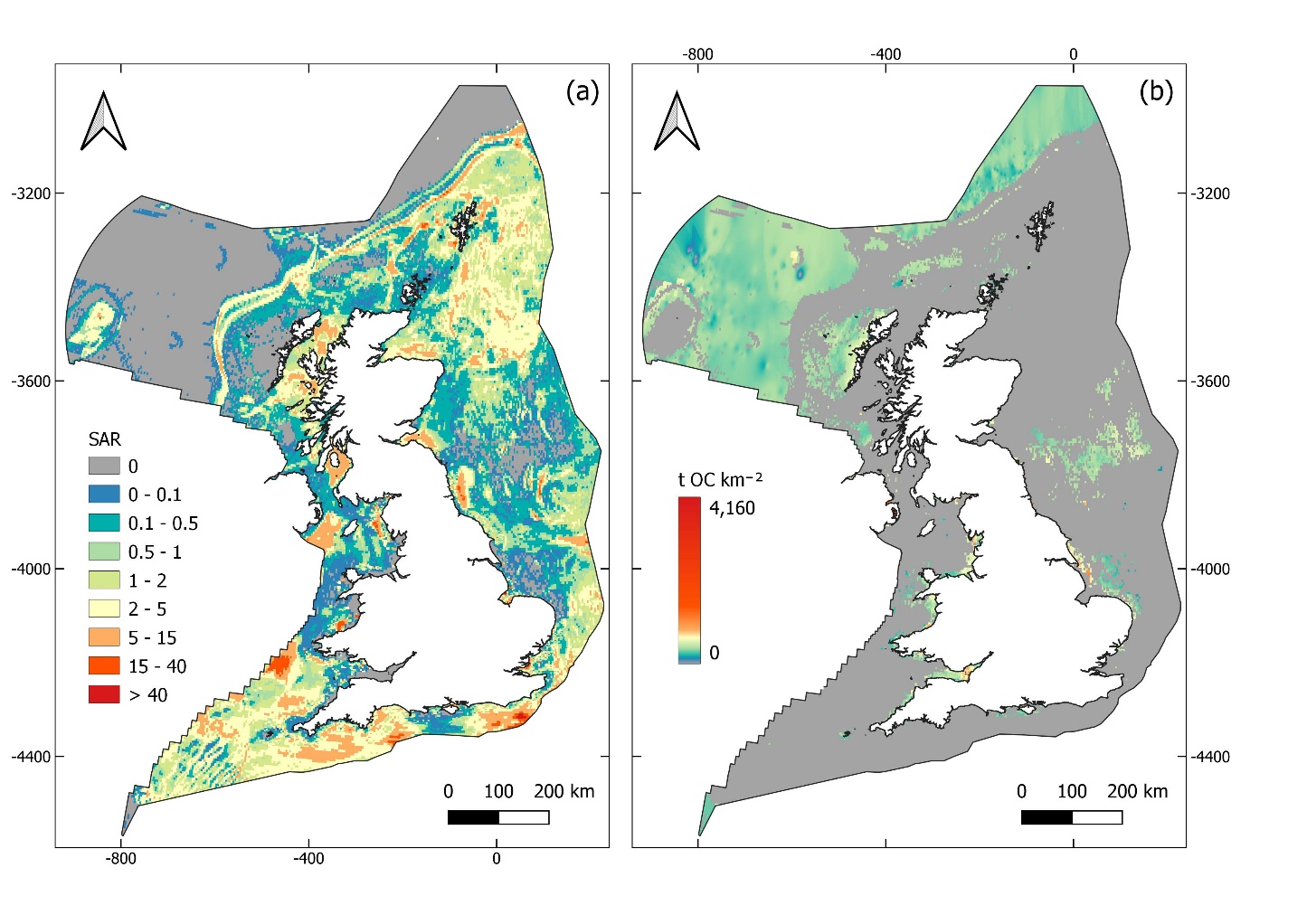

**Fig. S4.** Maps of the area and scale of mobile bottom fishing impact across the UK EEZ. (a) Fishing disturbance quantified by swept area ratio (SAR) of mobile bottom fishing gear use across study area based on data predominantly from the years 2009 to 2018. (b) Modelled concentration of organic carbon (OC) in unfished areas. Colours follow an exponential scale from low concentrations in blue to high concentrations in red, areas in grey indicate fished pixels.

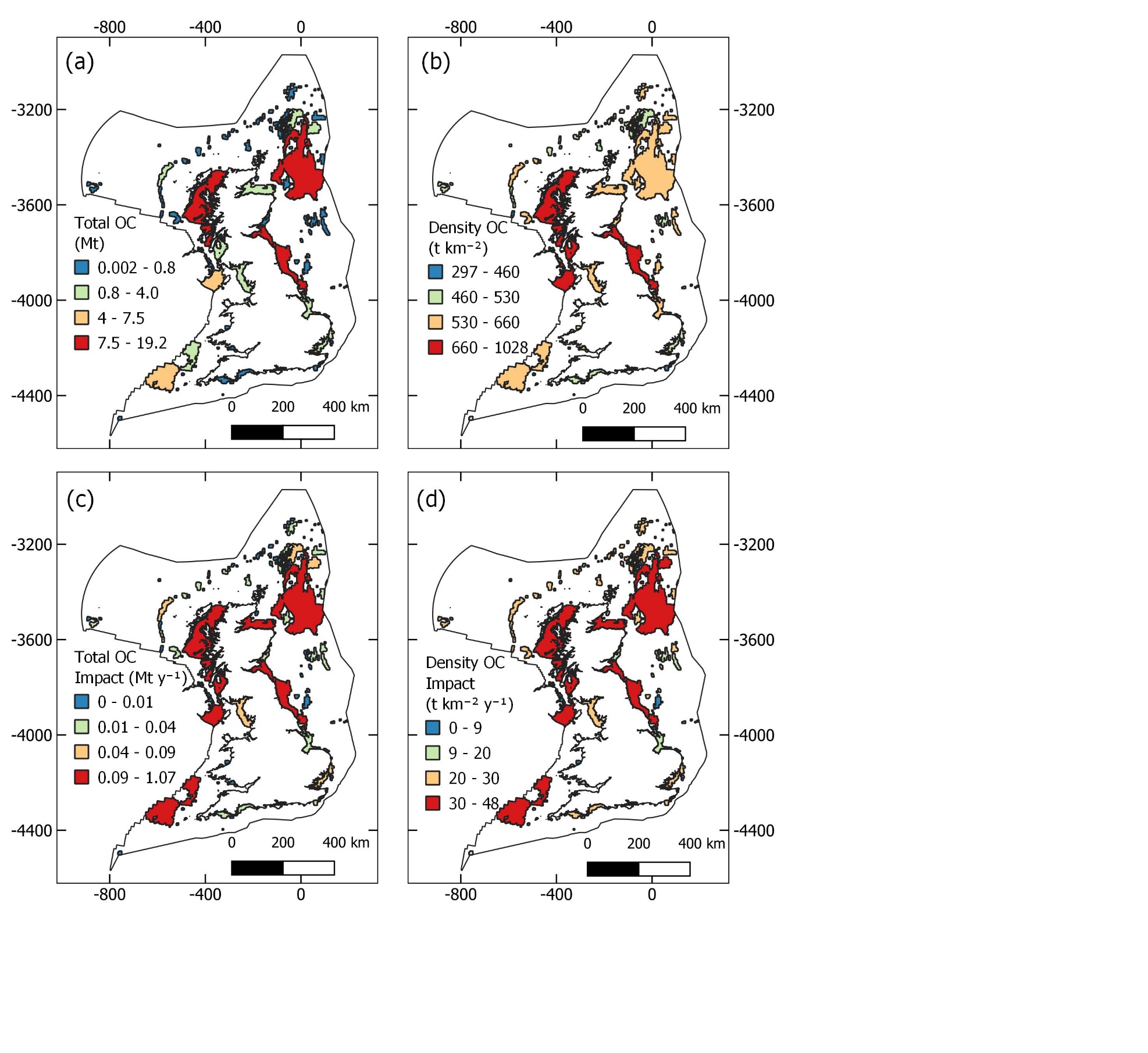

**Fig. S5** All selected seabed organic carbon priority areas. Sediment organic carbon (OC) priority areas were first selected based on identifying aggregations of fished pixels with high concentrations of predicted OC loss and/or OC stock. This lead to 207 areas being selected. All selected priority areas are shown with (a) the total OC stock within each area, (b) the density of OC within each area, (c) estimated total loss of OC from mobile demersal fishing and (d) the density of OC loss.

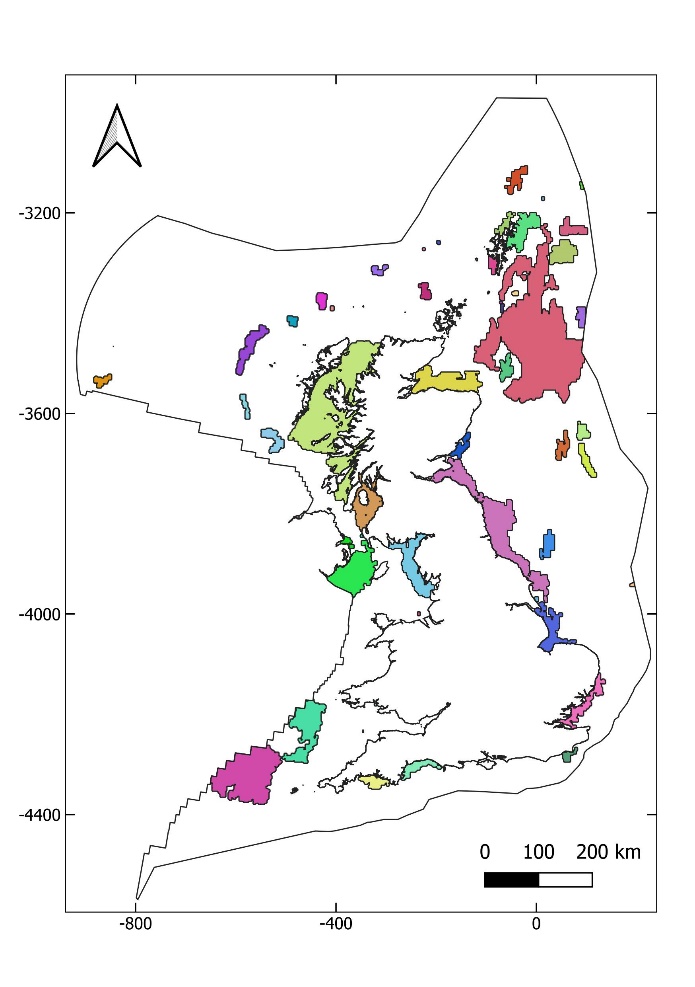

**Fig. S6** Potential seabed organic carbon (OC) priority areas. 55 potential priority areas were selected from those shown in Figure S4 by selecting those areas which met one of the following four criteria: 1) contained at least 0.1% of total predicted OC loss across the study site; 2) density of OC loss within the top 10% of all selected areas; 3) contained at least 0.1% of total OC stock across the study site; 4) density of OC stock within the top 10% of all selected areas. Colours only present to aid denotation of individual priority areas.

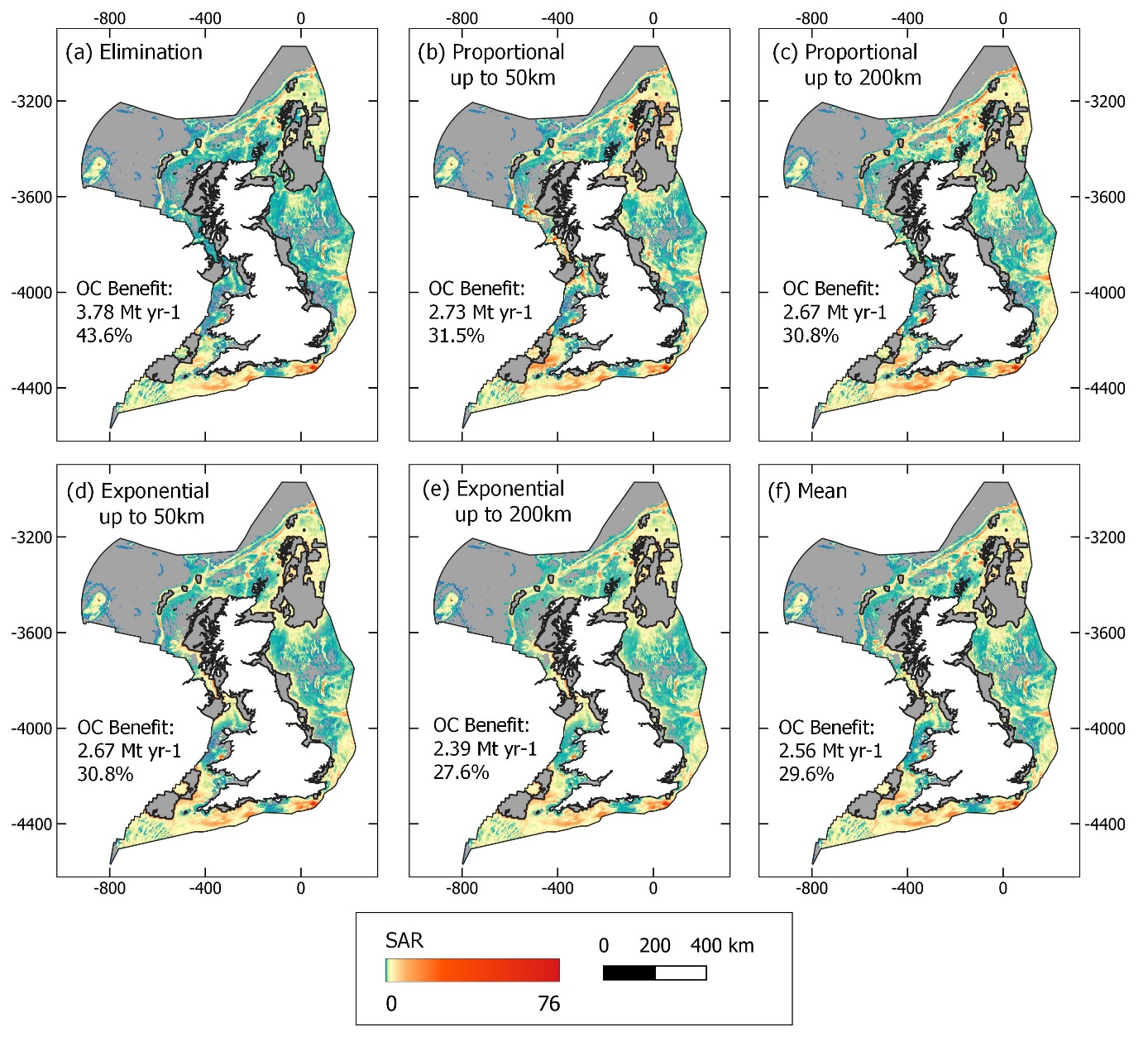

**Fig. S7.** Swept area ratio (SAR) following different fisheries displacement scenarios based on the establishment of mobile bottom gear closures in all priority areas. “Organic carbon (OC) benefit” reports the estimated saving in OC compared to baseline fishing disturbance (Fig. 2a, Fig 3a). Maps that show the resultant estimated loss of OC from each displacement scenario are shown in Fig. S7.

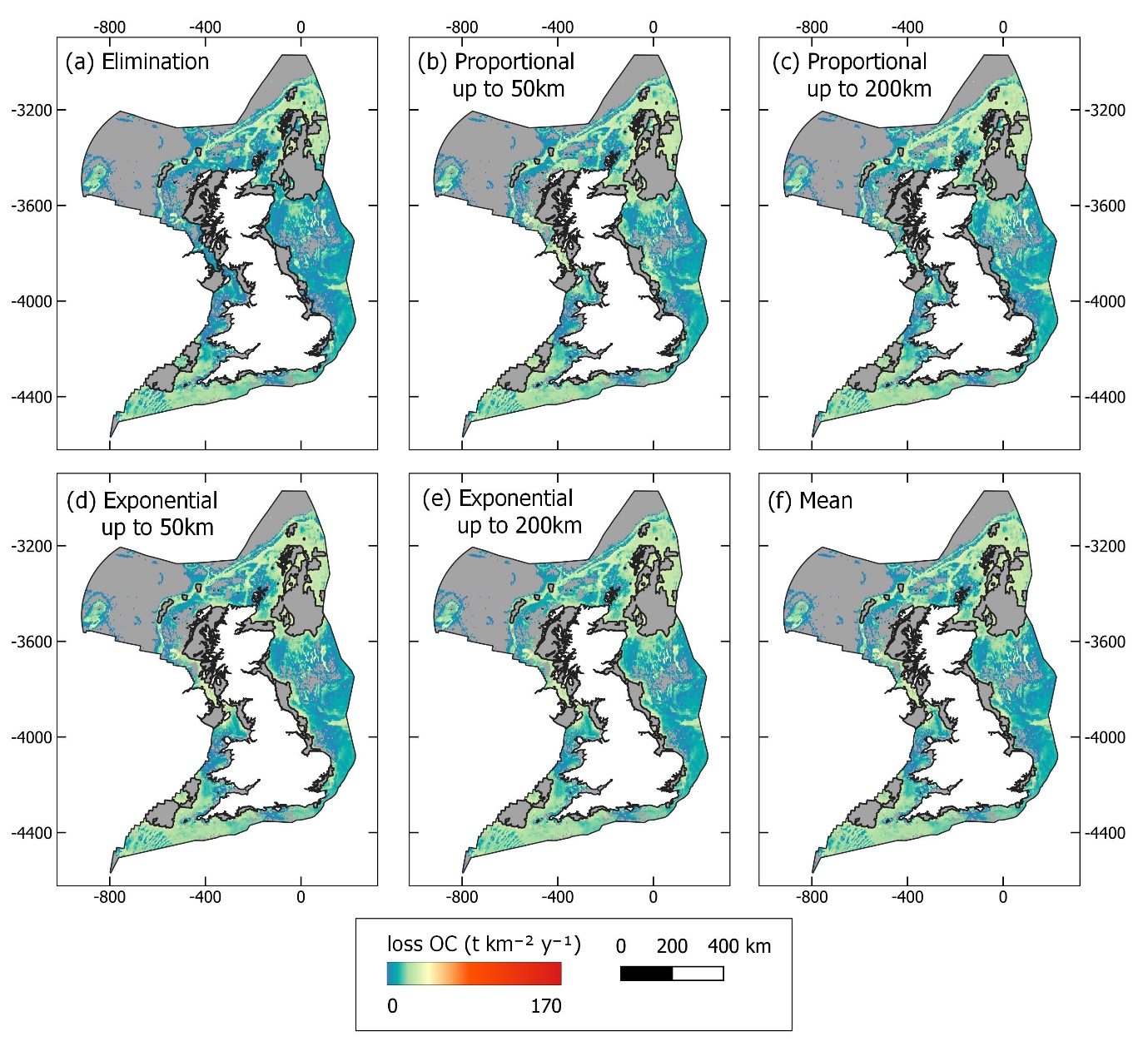

**Fig. S8.** Predicted loss of organic carbon (OC) due to modelled fisheries displacement from proposed blue carbon priority areas as shown in Figure S6.

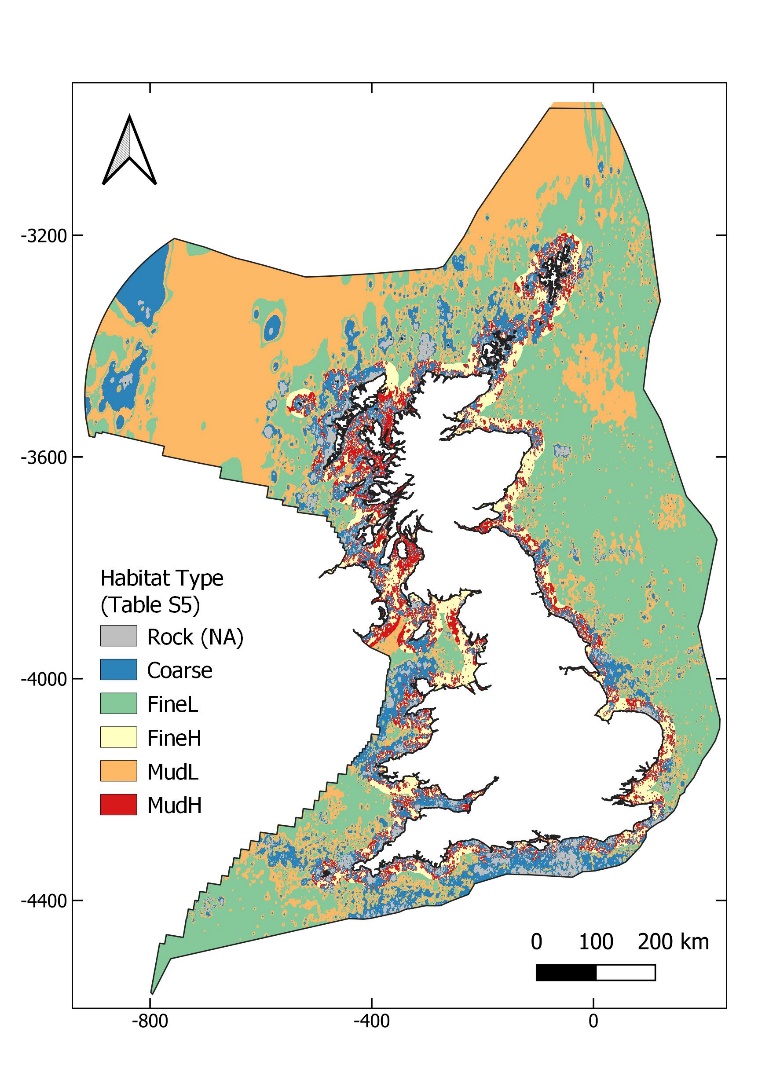

**Fig. S9** Habitat type assigned to each pixel across the study site to align with De Borger, Tiano, Braeckman, Rijnsdorp and Soetaert (*5*). Classification based on sediment type and from Smeaton, Hunt, Turrell and Austin (*6*) and distance from shore. The details of the classification are shown in Table S3.
